# Colocalization between bone loading and metabolic activity in the human tibia in response to acute exercise

**DOI:** 10.64898/2026.10.05.756847

**Authors:** Olivia L Bruce, Ananya Goyal, Yael Vainberg, Anthony A Gatti, Feliks Kogan

## Abstract

Bone strain and remodeling are important factors for bone health. Improved characterization of relationships between these factors in response to acute loading will improve our understanding of bone maintenance, adaptation to load, and pathophysiology. The purpose of this study was to evaluate colocalization between finite-element-estimated bone strain and [^18^F]sodium fluoride positron emission tomography imaging-estimated subchondral bone remodeling in the tibial diaphysis in response to acute exercise. We hypothesized that the exercise-induced change in standardized uptake values would be greatest in regions of higher strain. 26 participants underwent bilateral positron emission tomography/magnetic resonance scans before and after a bout of stair ascent and descent exercise. Finite element analysis was performed to calculate principal strain distributions in the tibial diaphysis. [^18^F]sodium fluoride standardized uptake values were calculated from pre- and post-exercise positron emission tomography data and mapped to the tibial surface. Mean pre-exercise and exercise-induced change in standardized uptake values were calculated and compared between regions defined by strain percentile. Both normalized mean pre-exercise and exercise-induced change in standardized uptake values increased with maximum principal strain in a pattern suggesting an exponential relationship. Pre-exercise and exercise-induced change in standardized uptake value were notably much greater in the region of highest maximum principal strains (estimated marginal means = 76.9 and 71.0 percentile, respectively) when compared to all other regions of the tibial diaphysis (≤ 61.1 and ≤ 57.0 percentile, respectively, p < 0.001). This study is the first to directly show a relationship between bone mechanical loads and acute [^18^F]sodium fluoride uptake in humans *in vivo*. Our results highlight the influence of tensile strain magnitude on bone metabolic activity in response to acute loading. Furthermore, we demonstrate the utility of a combined finite element analysis and positron emission tomography approach to interrogate relationships between bone loading and bone metabolic response to acute exercise.

**Lay summary:** This study aimed to understand the relationship between bone loading and bone metabolic activity, important factors influencing bone health. Before and after exercise, 26 participants underwent imaging following injection with [^18^F]sodium fluoride, a radiotracer that localizes to areas in bone with increased perfusion and/or increased remodeling activity. Tracer uptake was spatially compared to bone loading distribution, calculated using personalized models. Exercise-induced change in tracer uptake was increased in regions of high tensile strain, highlighting the importance of loading magnitude in bone metabolism. This combined imaging and modeling approach is a promising method to interrogate relationships between loading and bone metabolism.

## 1. Introduction

Bone strain and bone remodeling are interconnected factors associated with the maintenance of bone health. Muscle and joint forces applied to the bone interact with the bone’s structure and mechanical properties, resulting in bone strain (Edwards 2018). Bone strain, or some consequence thereof, is thought to stimulate targeted bone remodeling and adaptation (Allen & Burr 2014). Imbalances or abnormalities in this relationship impact the pathophysiology of bone diseases, leading to skeletal fragility and bone stress injuries (Hoenig et al., 2022; Recker et al 2004). Despite these clinical consequences, our current understanding of the bone’s metabolic response to acute loading is limited and is largely based on animal models (Li et al., 2005; Tomlinson et al., 2012; Silva et al., 2006; Uthgenannt & Silva, 2007).

Positron emission tomography (PET) is a promising molecular imaging technique that can assess bone metabolism in humans *in vivo*. [^18^F]-sodium fluoride ([^18^F]NaF) is an established PET radiotracer used clinically as a marker of bone turnover (Watanabe et al., 2015; Cronlein et al., 2015; Raynor et al., 2016; Rhodes et al., 2020). [^18^F]NaF begins to accumulate in the bone extravascular space and on the bound water shell of exposed hydroxyapatite crystals within seconds to minutes of injection (Blau et al., 1972). Thus, elevated [^18^F]NaF uptake in a region may be driven by an increase in exposed bone surface (e.g., microdamage or resorption canals) and/or increased bone blood flow (Genant et al. 1974; Goyal et al., 2024).

Recent studies have assessed bone metabolic changes in response to acute loading using [^18^F]NaF PET, primarily in subchondral bone in the knee joint (Watkins et al. 2023). Haddock et al. (2019) scanned the knees of 12 healthy participants before and after a step-up and drop-down exercise. Standardized uptake value (SUV), a measure of tracer uptake within individual participant that accounts for the decay of the tracer over time and the participant’s body mass, increased post-exercise by 47-131% in various knee subchondral bone subregions; pharmacokinetic modeling, which can estimate individual rates of bone blood flow and bone mineralization, indicated these increases were largely driven by an increase in bone perfusion. Similarly, Watkins et al. (2022) scanned 10 patients with knee osteoarthritis before and after a single-leg squat to fatigue exercise. [^18^F]NaF SUV, tracer extraction, and bone perfusion increased in the exercised knee when compared to the immobilized knee. Importantly, both studies observed side-to-side and regional variations in the SUV increases with exercise, which were hypothesized to reflect a load-dependent metabolic response to the exercises (Haddock et al., 2019; Watkins et al., 2022). However, neither study quantified the distribution of load in the bones to confirm this hypothesis. To address this gap, estimates of bone strain distribution, which can be achieved using participant-specific finite element modeling (Grassi et al., 2013; Haider et al., 2020), can be utilized. Thus, the purpose of this study was to evaluate colocalization between finite element-estimated bone strain and [^18^F]NaF uptake in the tibial diaphysis in response to acute loading. We hypothesized that regions experiencing higher finite element-estimated strain would exhibit a larger exercise-induced increase in bone metabolism as measured by the change in [^18^F]NaF PET SUV.

## 2. Materials and Methods

### 2.1 Participants and study design

Preliminary analysis of one individual fit a linear model to evaluate the node-wise relationship between maximum principal (tensile) strain and exercise-induced percent increase in SUV. Using the observed effect size (rho = 0.40) and assuming a two-tailed test and alpha = 0.05 for a point biserial model, the estimated sample size needed to achieve 80% power was 44 (G*Power v.3.1, Faul et al., 2009).

26 participants (n=52 tibiae) were recruited from the community. Participants were primarily recruited for another study. 11 participants had self-reported unilateral knee pain and the other 14 were asymptomatic. Participants had no conditions or pain affecting the tibia, besides knee pain. The study was approved by the university’s Institutional Review Board, and all participants gave written informed consent prior to participating in the study. Participants were asked to abstain from weight-bearing exercise for the 24 hours prior to their scan appointment.

Participants received an intravenous injection of 93 ± 4.4 MBq of [^18^F]NaF. Following a waiting period of approximately 30-35 minutes (while the knee was scanned for another study) bilateral PET and MR images were simultaneously acquired using a 3T whole-body time-of-flight hybrid PET/MRI system (GE SIGNA, GE Healthcare, Milwaukee, WI). Following this first scan, participants ascended and descended a flight of stairs eight times (135 steps up and down, self-paced). Participants were then repositioned within the PET/MRI scanner, received a second injection of 93 ± 4.4 MBq of [^18^F]NaF, and scanned using the same imaging protocol as the first scan.

### 2.2 Image acquisition details

Three overlapping segments (3 PET beds) were used to cover the entire tibia with 4 minutes of data acquisition in each PET bed position. Both legs were scanned with the following PET parameters: list-mode, field of view = 600 mm (transaxial), 250 mm (axial), reconstructed matrix = 384 x 384, reconstructed slice thickness = 2.78. 3D Axial T1-wieghted MR images of the lower legs were acquired in three overlapping sections using a spoiled gradient echo (SPGR) sequence with TR = 11.94 ms, TE = 5.77 ms, field of view 320 mm, matrix = 1024 x 1024, slice thickness = 5.00 mm.

### 2.3 Image analysis

#### 2.3.1 Finite element models

The T1-weighted images were cropped and split into left and right sides. The three overlapping sections for each lower leg were then stitched together. The left and right tibia and fibula were manually segmented from the post-exercise MR images and then meshed (edge lengths = 0.75-1.75 mm, mesh grading = 0.3, number of elements: tibia = 165186 – 260669, fibula = 23779 – 45759) using NGSolve, v6.2.2304 (https://ngsolve.org). A prior convergence analysis observed less than 4% change in peak strain when the number of elements were approximately doubled from 132740 to 270002 for tibia, and 20241 to 41077 for fibula (Bruce et al., 2022).

Image signal intensity values were assigned to each element in the meshes based on the underlying MR image in Bonemat (v3.2; Taddei, 2007). Elements with grayvalues below 25% of the maximum signal intensity in the mesh were labelled as cortical bone. The 25% threshold was chosen based on an assessment of a frequency plot of normalized gray values within the tibia mesh for a randomly chosen example participant. In addition, the root mean square error of diaphysis surface maximum and minimum principal strains between finite element models using 20% and 25% thresholds was minimal (24.9 – 41.4 µε, <5%). All elements with a vertex on the surface of the mesh were also labelled as cortical bone. Non-cortical elements in the proximal and distal 15% of bone length were labelled as trabecular bone. All remaining elements were labelled as bone marrow. Generic linear elastic orthotropic material properties (E: Young’s modulus, G: Shear modulus, and v: Poisson’s ratio) were assigned for cortical bone (E_1_ = 11.70 GPa, E_2_ = 12.20 GPa, E_3_ = 20.70 GPa, G_12_=4.10 GPa, G_13_ = 5.17 GPa, G_23_ = 5.70 GPa, v_12_ = 0.420, v_13_ = 2.37, v_23_ = 0.231; Rho et al., 1996). Isotropic material properties were assigned for trabecular bone (E = 0.40 GPa, v = 0.3) and bone marrow (E = 0.30 GPa, v = 0.45; Tarapoom & Puttapitkporn 2016).

Generic knee contact forces and moments at the time of peak resultant knee contact force during stair descent were calculated for five participants in the Orthoload database (file list in Supplementary Information; Bergmann G. and Damm P., 2008). The peak knee contact forces and moments were normalized to body mass, then averaged across participants. The generic normalized mean knee contact forces and moments were scaled by each participant’s body mass and applied as concentrated forces/moments at the intercondylar eminence.

Boundary conditions were similar to those reported in Haider et al (2020). Briefly, one point on the intercondylar eminence was constrained to allow only axial motion and was coupled (continuum distributing) to the tibial plateau surface. A pinned constraint was used at the ankle to allow only sagittal- and coronal-plane rotation at the ankle joint: a virtual ankle centre of rotation was defined as the midpoint between the lateral and medial malleoli, constrained to disallow translation, and coupled (continuum distributing) to the surfaces of the talotibial and talofibular surfaces. One point on the medial malleolus was fixed in the anterior-posterior direction. The distal fibula was rigidly tied to the distal tibia. At the proximal tibiofibular joint, springs with stiffness of 133 N/mm and 109 N/mm were modeled to approximate the anterior and posterior ligaments, respectively (Marchetti, 2017). Furthermore, the tibiofibular joint was constrained to prevent an opening perpendicular to the joint surface. Finite element models were evaluated in Abaqus (v2024, Dassault Systemes, France). Maximum (tensile) and minimum (compressive) principal strains at tibial surface nodes were extracted.

#### 2.3.2 SUV: Standardized uptake values (SUV) were calculated for pre- and post-exercise PET images using the equation

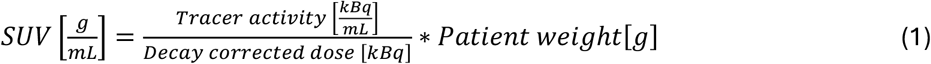

SUV is a measure of [^18^F]NaF concentration normalized to decay-corrected quantity of the injected [^18^F]NaF and the patient’s body mass (Goyal et al., 2024). The decay-corrected dose from each injection was calculated using the equation:

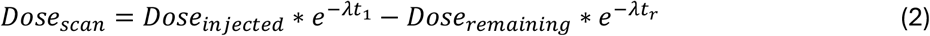

where *Dose*_*injected*_ is the injected dose of tracer, *Dose*_*remaining*_ is the amount of the dose remaining in the syringe following injection, *λ* is the decay constant of the radioactive nucleus, *t*_*1*_ is the time from dose injection to the time of the scan, and *t*_*r*_ is the time from dose injection to the time of measurement of the remaining dose in the syringe. For post-exercise images, the amount of circulating tracer from both the first and second injections was accounted for in the dose calculation. The decay-corrected amount of circulating tracer from the first injection at the time of the second scan was calculated using Equation 2, but with t1 representing the time between the first injection and the second scan. These two doses were then summed to determine the total decay-corrected dose for the post-exercise scan.

The cropped and stitched T1-weighted anatomic MR images were used for registration between pre- and post-exercise scans. For each left or right leg, the post-exercise stitched image volume of the isolated lower leg was rigidly registered to the corresponding pre-exercise image volume using general intensity-based methods (Johnson et al., 2007). In cases where the lower leg was translated by more than ∼1 cm or rotated more than ∼10 degrees, manual alignment was performed as a first step prior to automated rigid registration. The resulting rotation matrix was used to align the SUV images. The exercise-induced change in SUV was then computed by subtracting the pre-exercise image volume from the post-exercise image volume. Pre-exercise and exercise-induced change in SUV values were then mapped to the tibia mesh surface. For each node on the tibial surface, the maximum SUV or change in SUV ± 10 mm along a line normal to the surface was determined.

### 2.4 Statistics

Tibial diaphysis surface nodes (defined as 20-80% of tibia length) were binned into ten pointsets by strain percentile (0-10%, 11-20%,… 91-100%; Figure 1). For minimum principal strain, the absolute value was used. Pre-exercise SUV and exercise-induced change in SUV were normalized to corresponding range within each tibial diaphysis, expressed as percentile. Mean normalized SUV parameters were then calculated for each strain pointset.

**Figure 1:**
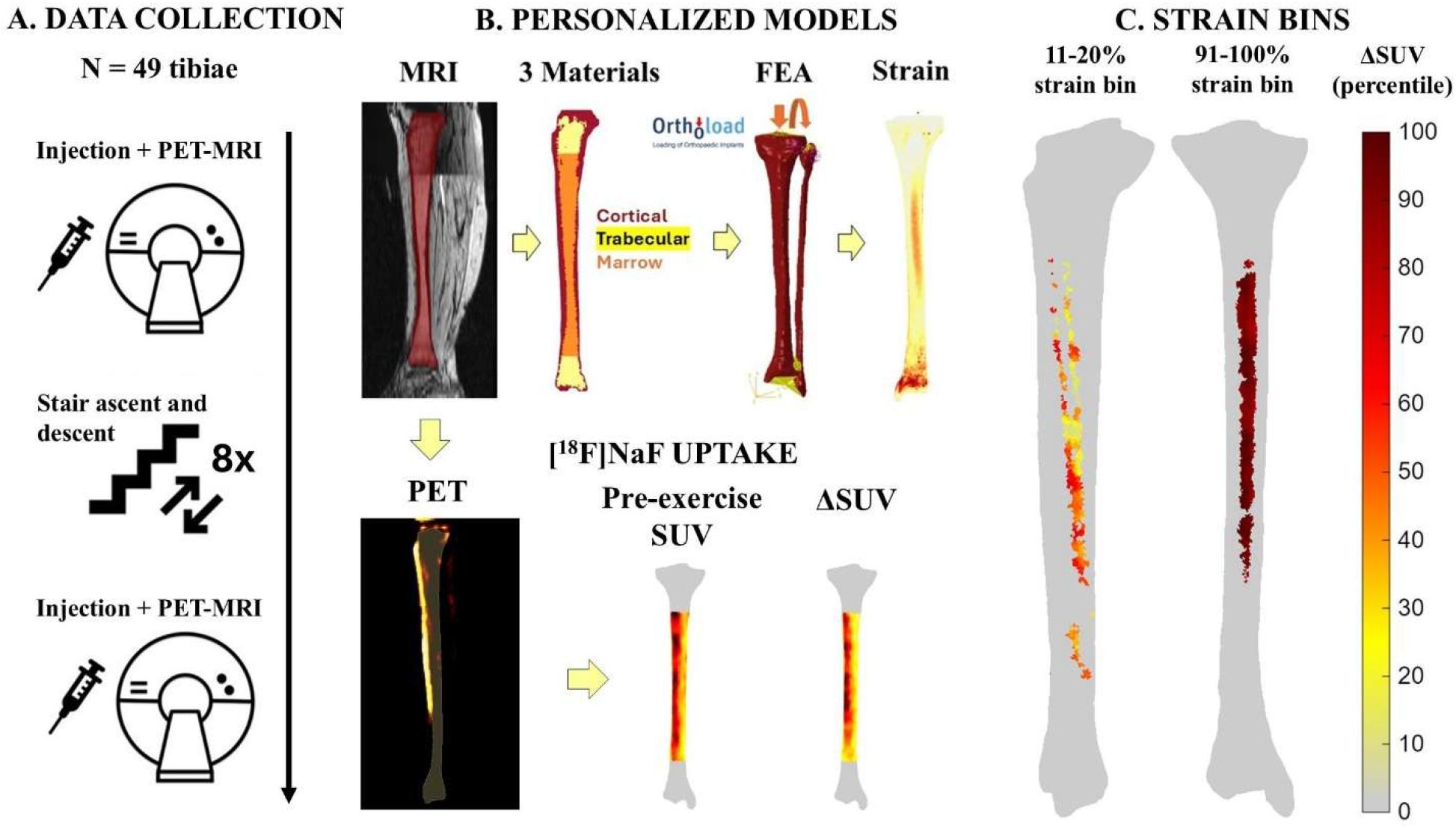
Overview of methodological workflow. Participants were injected with [^18^F]NaF and then underwent simultaneous PET-MRI scanning. The participants performed a stair climbing exercise, then repeated the injection and scanning protocol. MRI images were used to generate finite element models of the tibia and fibula to calculate bone strain, and for image registration. Pre-exercise and exercise-induced change in standardized uptake values of [^18^F]NaF were calculated using the PET images and were mapped to the surface of the tibial diaphysis. Mean SUV was calculated in regions defined by strain percentile within each tibia.

Statistical analyses were performed in R (v4.5.0). Linear mixed effects models were used to evaluate colocalization between strain and SUV. Strain bin, knee pain (defined as: affected leg, contralateral leg, healthy control), and an interaction term (strain bin x knee pain) were modeled as categorical fixed effects. A random intercept for participant was included in the model to account for variance in participant baselines. QQ plots of model residuals were visually assessed for normality.

Mean exercise-induced change in SUV for maximum principal strain bins was natural log transformed to adjust for a right-skewed distribution and non-normal model residuals. Pairwise comparisons were performed using the *emmeans* package (v1.11.0, Lenth 2025). p-values were adjusted for multiple comparisons using Tukey’s method.

## 3. Results

### 3.1 Participant demographics

One participant was excluded following data collection due to a benign lesion in their lower leg; the remaining 25 participants were included in the analysis (18 female, 7 male, 45.0 ± 17.5 years, 1.70 ± 0.09 m, 45.0 ± 17.5 kg). The right leg of one of the participants was not entirely captured in the MR image volume and was excluded, resulting in a total of 49 tibiae.

### 3.2 SUV and strain patterns

The highest SUV and exercise-induced changes in SUV measurements typically occurred along the proximal two-thirds of the anterior tibial crest (Figure 2). Within each tibia, the exact patterns varied between pre-exercise SUV and exercised-induced change in SUV (Figure 2). Hotspots, or areas of focal PET uptake, occurred on the posterior surface as well in approximately 50% of the tibiae (24 of 49 for pre-exercise SUV and 27 of 49 for change in SUV). While the highest maximum principal (tensile) strains were typically observed along the anterior crest, in contrast, the highest minimum principal (compressive) strains were observed on the posterior surface of the tibial diaphysis.

**Figure 2:**
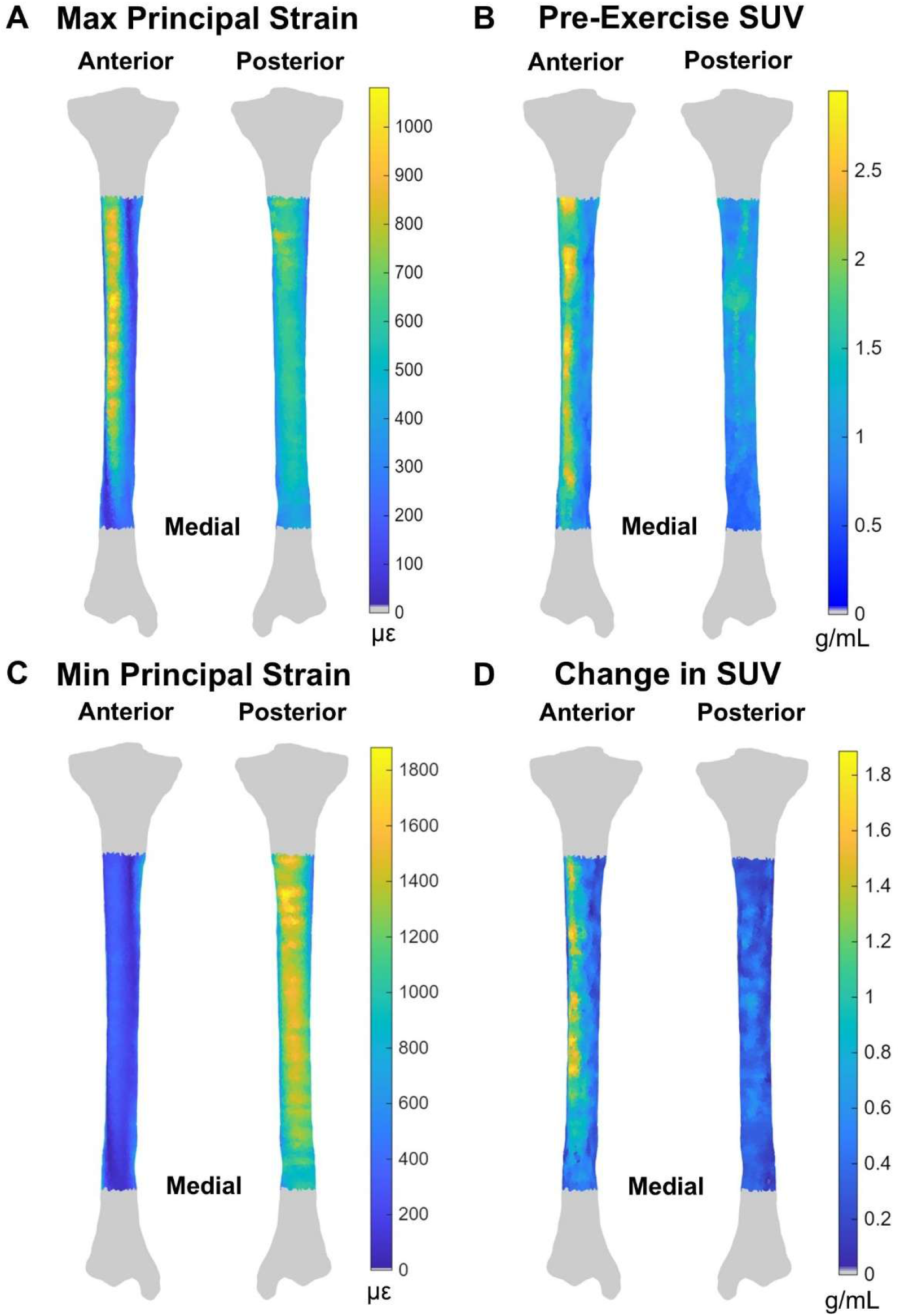
Examples of strain and SUV maps across the tibial diaphysis for one representative participant. A left tibia is shown.

### 3.3 Knee pain

There were no interactions between strain bin and pain for any SUV parameters (p > 0.05). There were also no differences between limbs or between unilateral knee pain and healthy participants for any SUV parameters (p > 0.05).

### 3.4 Colocalization

Mean normalized pre-exercise SUV increased with maximum principal strain percentile, beginning at the 41-50^th^ percentile strain region (Figure 3A, detailed pairwise comparisons shown in Figure 3C, p ≤ 0.04). The largest increase in pre-exercise SUV from one strain percentile region to the next was observed between the two highest strain percentile regions. Estimated marginal means for normalized pre-exercise SUV significantly increased from 61.1 percentile in the 81^st^-90^th^ percentile strain region to 76.9 percentile in the 91^st^-100^th^ percentile strain region (p<0.001). Similarly, mean normalized exercise-induced change in SUV increased with strain percentile beginning at the 51-60^th^ percentile strain region (Figure 3B, detailed pairwise comparisons shown in Figure 3D, p ≤ 0.03). Again, there was a marked increase in exercise-induced change in SUV from the 81^st^-90^th^ percentile strain region (estimated marginal mean = 57.0 percentile) to the 91^st^-100^th^ percentile strain region (estimated marginal mean = 71.0 percentile, p < 0.001).

**Figure 3:**
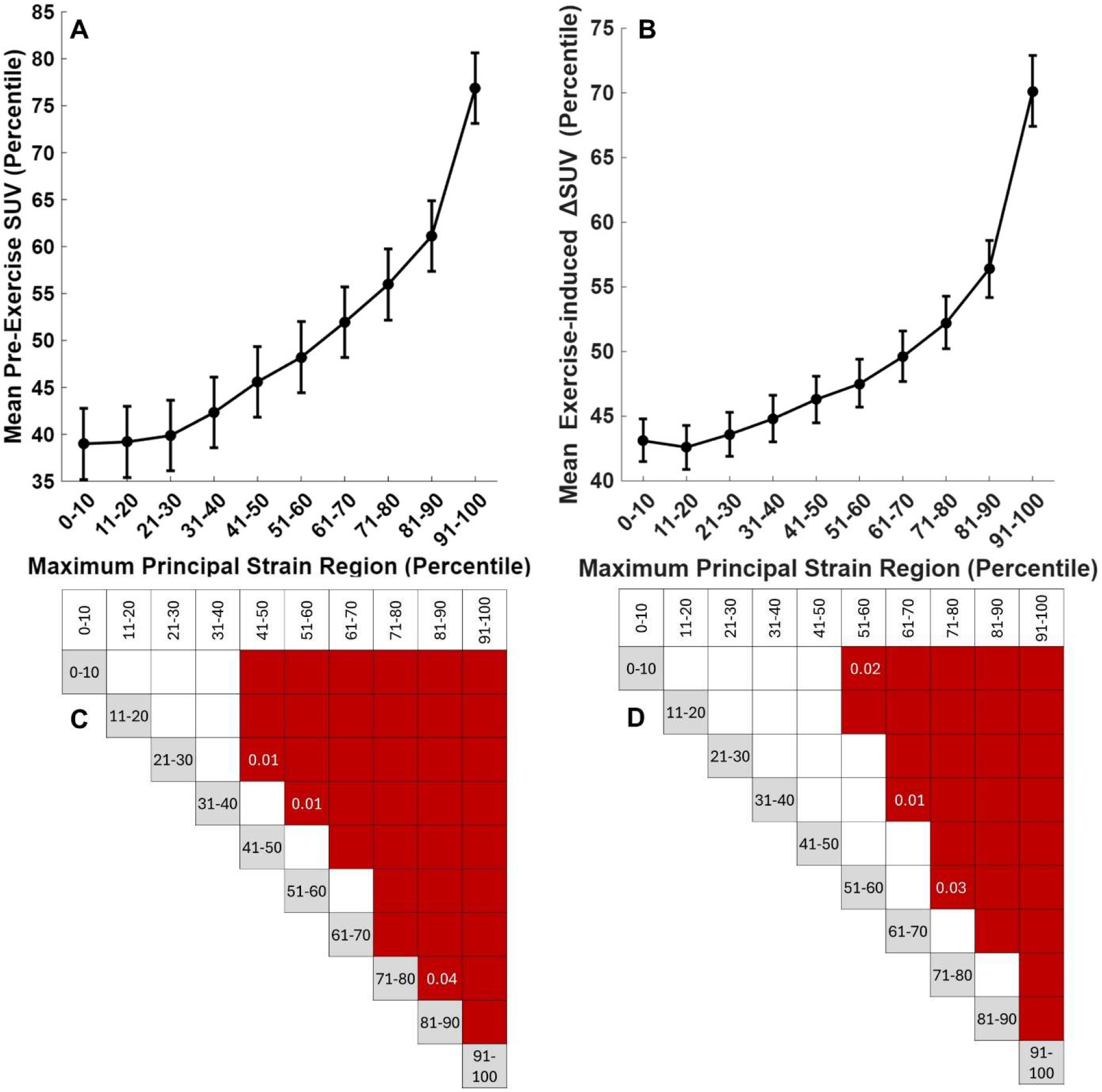
Estimated marginal mean and standard error (normalized) SUV across strain bins. Top: A) Mean SUV and B) exercise-induced change in SUV demonstrated an exponential relationship with tensile (maximum principal) strain. There is little change in lower strain regions and a marked increase at the greatest strain bin. Bottom: Corresponding charts of pairwise comparisons of SUV between strain percentile bins (x- and y-axes). Significant comparisons are shaded red. p-values were < 0.01 unless otherwise shown in each red square.

In contrast, mean normalized pre-exercise SUV and exercise-induced change in SUV did not demonstrate a directional relationship, instead illustrating a sinusoidal pattern. SUV variables were statistically greater in the low minimum principal strain regions corresponding to the anterior crest when compared to higher minimum principal strain regions corresponding to the posterior surface (Figure 4A and 4B, detailed pairwise comparisons shown in Figure 4C and 4D, p ≤ 0.047). Mean normalized pre-exercise SUV was greater in the 21^st^-50^th^ percentile minimum principal strain regions (estimated marginal means = 55.1 – 64.1 percentile) when compared to the 51^st^-80^th^ percentile minimum principal strain regions (estimated marginal means = 39.3 – 45.5 percentile, p < 0.001). Similarly, mean normalized exercise-induced change in SUV was greater in the 21^st^-50^th^ percentile minimum principal strain regions (estimated marginal means = 56.5 – 63.4 percentile) when compared to the 51^st^-90^th^ percentile minimum principal strain regions (estimated marginal means = 39.6 – 50.2 percentile, p < 0.001).

**Figure 4:**
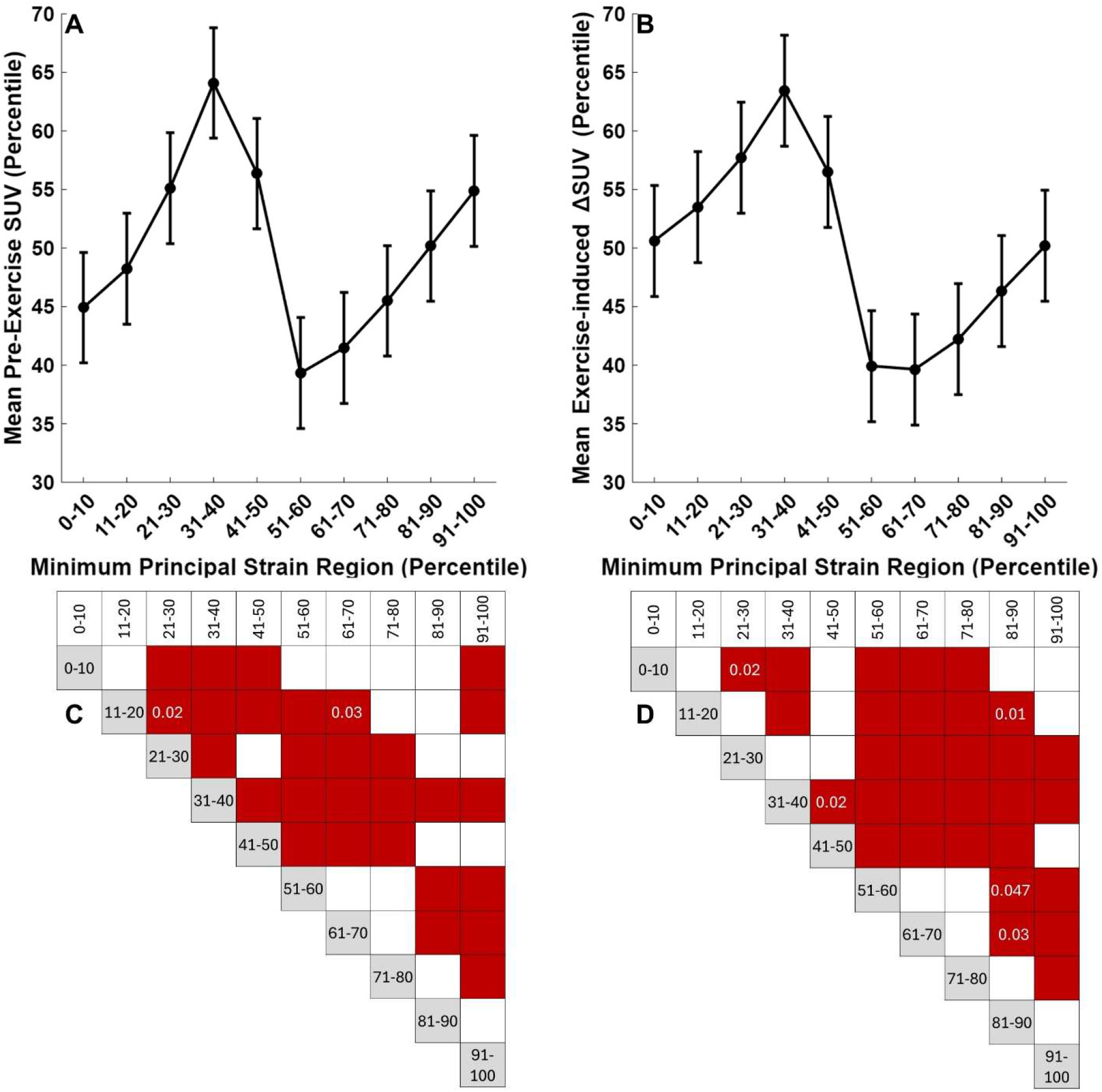
Estimated marginal mean and standard error (normalized) SUV across strain bins. Top: Mean pre-exercise SUV and exercise-induced change in SUV did not show a directional relationship with compressive (minimum principal) strain. Bottom: Corresponding charts of pairwise comparisons of SUV between strain percentile bins (x and y axis). Significant comparisons are shaded red. p-values were < 0.01 unless otherwise shown in each red square.

## 4. Discussion

The purpose of this study was to evaluate colocalization between finite element model-estimated bone strain and [^18^F]NaF PET imaging-estimated bone metabolic activity in the tibial diaphysis in response to acute loading. Both pre-exercise SUV and exercise-induced change in SUV in the tibial diaphysis were typically greatest along the proximal two-thirds of the anterior crest. SUV parameters increased with maximum principal (tensile) strain and were particularly elevated in the regions of the tibial diaphysis experiencing the highest tensile strains, suggesting an exponential relationship. Meanwhile, there was no clear directional relationship between SUV parameters and minimum principal (compressive) strain. The results correspond with findings from animal models (Li et al., 2005; Tomlinson et al., 2012; Silva et al., 2006; Uthgenannt & Silva, 2007) and demonstrate the potential utility of this dual-pronged approach to interrogate bone strain-metabolism relationships to better understand bone adaptation, pathophysiology, and the acute effects of biomechanical interventions.

Both pre-exercise SUV and change in SUV following stair ascent and descent exercise demonstrated a pattern across participants whereby the greatest SUV was typically observed along the proximal two-thirds of the anterior crest. The pre-exercise SUV pattern may be indicative of small local increases in perfusion, vascularity, and remodeling activity associated with the participants’ day-to-day loading in the week preceding the experiment (Muir et al., 2007, Silva et al. 2006). The exercise-induced change in SUV illustrated a similar pattern. The greatest changes were observed along the anterior crest in the middle third of the diaphysis and more proximally near the tibial tuberosity where loads from bending and the patellar ligament insertion, respectively, are expected to be high. Similarly, when hotspots, areas of focal increase in PET signal, were observed on the posterior surface, they often occurred at the mid-diaphysis where loads are expected to be high. The local increase in SUV we observed in response to acute exercise is consistent with previous studies of [^18^F]NaF uptake in the knee following exercise and likely indicates local increases in bone perfusion and blood flow (Haddock et al., 2019; Watkins et al., 2022). In these previous studies, the greatest exercise-induced change in SUV occurred in the patella and the tibial tuberosity. Furthermore, the change was greater in the knee that performed a step-up when compared to the knee that performed a step down (Haddock et al., 2019). This result is consistent with what would be expected based on general assumptions of mechanical loading distributions in the knee, where the patella and tibial tuberosity are subject to high muscle forces transmitted through the patellar tendon and ligament. Importantly, however, this was an observational finding as load was not quantified.

Here, we expanded on the previous studies to directly evaluate the relationships between mechanical load and exercise-induced changes in SUV. We hypothesized that change in [^18^F]NaF uptake following acute exercise would be greatest in areas of high strain. The results support our hypothesis, demonstrating that the areas of greater change in SUV following exercise do indeed colocalize with regions of the highest finite element-estimated maximum principal strains within the tibial diaphysis. This pattern is consistent with current understanding of targeted remodeling and highlights the importance of strain magnitude as a factor affecting this process. Bone microdamage is proportional to the number of loading cycles multiplied by the load (stress or strain) raised to a power (Edwards, 2018), echoing the exponential pattern we observed. This microdamage has been associated with increased perfusion and blood flow and is thought to be one of the main factors triggering a targeted bone remodeling response (Allen & Burr, 2019; Muir et al., 2007).

In contrast, there was no apparent directional relationship between SUV parameters and minimum principal strain, which is highest on the posterior surface of the diaphysis. The lack of [^18^F]NaF uptake in areas of highest compressive strain is likely due to the difference in bone stiffness and strength between compression and tension. Damage initiates at lower strain magnitudes and accumulates faster in bone loaded in tension when compared to compression (Pattin et al., 1996; Burr et al., 1998). Furthermore, maximum principal strains have demonstrated stronger relationships with fatigue life in whole bone loading tests when compared to minimum principal strains (Haider et al., 2021). Stair ascent and descent, a common activity of daily living, may not have resulted in sufficient stimulus on the compressive surface to induce a response. Further work testing higher magnitude loads is needed to evaluate this hypothesis. In addition, testing movements with greater loading magnitudes will allow improved characterization of the dose-response between SUV and tensile load magnitude.

A limitation of this study is that the finite element models were not fully personalized; while models were generated from participant-specific bone geometry data, bone material property distributions and muscle and joint contact forces were not available. Bone material properties in the tibia vary to a smaller degree and have a smaller effect on bone strains within and between healthy adult participants when compared to geometry (Bruce et al., 2022). As strain data were normalized to percentile, we do not expect that incorporating bone material property data would significantly change the results of this study. Generic knee joint contact forces for stair descent, computed from telemetric knee implant data and scaled by body mass, were also used (Bergmann & Damm, 2008). Incorporating participant-specific muscle and joint contact forces could influence the strain distribution pattern across the tibia. We expect that improving personalization would further strengthen the colocalization between finite element-estimated strain and SUV.

In this study, we used a novel approach combining [^18^F]NaF PET-MRI and finite element modeling to evaluate relationships between bone loading and metabolic activity in response to acute exercise in humans *in vivo*. Our results showed strong colocalization between high tensile strains and elevated [^18^F]NaF uptake, which aligns well with our current understanding of the bone’s response to mechanical stimuli. This is a feasible approach that could be used to better understand the role of loading in the pathophysiology of overuse injuries and other bone conditions. For example, bone stress injuries are thought to be a result of damage accumulation from steep increases in repetitive high magnitude loading and this process may be accelerated by the remodeling response, which would transiently weaken the bone through increased porosity (Hoenig et al., 2022). While structural changes in the cortical bone cannot be detected with computed tomography and MRI except at the highest severity grades (Fredericson et al., 1994), [^18^F]NaF PET imaging may be sensitive to local hotspots of elevated bone metabolic activity at lower grades (i.e., at an earlier stage in the development of the injury). Furthermore, the combination of finite element modeling and [^18^F]NaF PET imaging may be useful to evaluate the acute effects of biomechanical interventions in populations at increased risk of developing bone conditions such as bone stress injuries and osteoarthritis.

## Acknowledgements

Stanford Wu Tsai Human Performance Alliance (pre-doctoral and postdoctoral fellowships: OLB, AG, and AAG), Stanford DARE Graduate Fellowship (AG), NIH R01079431

The authors would like to thank Dawn Holley, Andrew Dreisbach, and Kim Halbert for their assistance with PET-MRI scanning.

## CRediT Statement

**Olivia L Bruce:** Conceptualization, Methodology, Formal analysis, Visualization, Writing – Original draft. **Ananya Goyal:** Conceptualization, Methodology, Investigation, Writing – Review & editing. **Yael Vainberg:** Investigation, Writing – Review & editing. **Anthony A Gatti:** Conceptualization,

Methodology, Writing – Review & editing. **Feliks Kogan:** Conceptualization, Methodology, Writing – Review & editing, Resources and supervision.

## Notes

**Conflict of Interest Disclosures:** AAG is a shareholder of NeuralSeg Ltd., NodeAI Diagnostics, and GeminiOV. All other authors have no conflicts of interest to disclose.

### Competing Interest Statement

AAG is a shareholder of NeuralSeg Ltd., NodeAI Diagnostics, and GeminiOV. All other authors have no conflicts of interest to disclose.

