## Supplementary Information for "Colocalization between bone loading and metabolic activity in the human tibia in response to acute exercise"

List of files from the Orthoload database (https://orthoload.com/database/) used to compute mean knee contact forces and moments at the time of peak resultant knee contact force during stair descent.

k1l_110108_2_37p

k2l_301107_1_144p

k3r_250309_1_105p

k4r_201108_2_3p

k5r_060809_1_56p

Bergmann G. and Damm P. (ed.), Julius Wolff Institute, Berlin Institute of Health at Charité Universitätsmedizin Berlin (2008) “OrthoLoad”.
